# Effects of lysine and methionine on mRNA expression of candidate transcription factors by primary bovine mammary epithelial cells

**DOI:** 10.1101/2024.06.01.596926

**Authors:** Boning Li, Ashlin M. Edick, Madison K. Fox, John Doelman, Sergio A. Burgos, John P. Cant

## Abstract

It has been established that essential amino acids (EAA) regulate protein synthesis in mammary epithelial cells by rapidly altering the phosphorylation state of translation factors. However, the long-term transcriptional response to EAA supply has been investigated much less. Eight transcription factors were selected as candidate mediators of EAA effects on mammary cell function via the amino acid response (*ATF4*, *ATF6*), mitogen-activated protein kinase (*JUN*, *FOS*, *EGR1*), and mechanistic target of rapamycin complex 1 (*MYC*, *HIF1A*, *SREBF1*). The objective was to determine if and when expression of these candidate genes was affected in primary cultures of bovine mammary epithelial cells more than 24 h after imposing an EAA deficiency, and to evaluate effects of EAA deficiency on protein synthesis, endoplasmic reticulum size, cell proliferation, and lipogenesis. Differentiated cells were cultured in 1 of 3 treatment media representing normal physiological concentrations of all amino acids (CTL), low lysine (LK), or low methionine (LM) for 24, 40, 48, or 60 h. Both LK and LM suppressed protein synthesis and activated *ATF4* expression, indicating the classic amino acid response pathway had been triggered. However, there was no effect of LK or LM on endoplasmic reticulum size, possibly related to elevated *ATF6* expression on LM. Expression of early response genes *JUN*, *FOS*, *EGR1* and *MYC* was not elevated by EAA deficiency but LM decreased *EGR1* expression. LM also increased expression of *HIF1A*. The *EGR1* and *HIF1A* expression results are consistent with the decrease in cell proliferation rate observed. Variable responses in *SREBF1* expression to LK and LM at different timepoints may have contributed to a lack of effect on lipogenesis rates. These findings indicate that EAA deficiency may inhibit mammary protein synthesis and cell proliferation through transcription factors.

## Introduction

Lysine (Lys) and methionine (Met) are essential amino acids (EAA) often found to be in deficient supply for maximal milk protein synthesis in the mammary glands of lactating dairy cows (1). It is well-documented that EAA can regulate milk protein yield by altering the phosphorylation state of translation factors through mechanistic target of rapamycin complex 1 (mTORC1), integrated stress response (ISR), and glycogen synthase kinase-3 signaling (2–5). This translational response does not appear to persist for long periods (days and weeks) over which dietary effects on milk protein yields are sustained and may just be part of the initial response (4,6). In addition to rapid effects on mRNA translation, there may be changes in gene transcription that contribute to the long-term effects of EAA on activities of the mammary epithelial cell. However, the transcriptional response has been investigated much less.

The ISR is the name given to the group of reactions induced by phosphorylation of the α subunit of eukaryotic initiation factor 2 (eIF2α). Four different eIF2α kinases sense a diverse array of nutritional, endoplasmic reticulum (ER), viral, and redox stresses and cause a decrease in global mRNA translation but an increase in translation of the mRNA for activating transcription factor 4 (*ATF4*) (7). *ATF4* belongs to the basic leucine-zipper family of transcription factors, including *ATF6*, *FOS*, and *JUN*, with which it homo- or hetero-dimerizes to induce expression of amino acid transporters, mRNA translation factors, and ER-resident proteins in what has been called the amino acid response pathway (AARP) (8). *ATF6* is itself an *ATF4* target (9) whose gene product can be activated by the ISR (10). An alternative to the *ATF4* pathway for transcription factor activation by EAA deficiency involves RAS/RAF/MEK/ERK signaling that stimulates the expression of *FOS*, *JUN*, and *EGR1* (11–13). Furthermore, mTORC1 inactivation by low EAA concentrations can decrease the expression of several genes encoding transcription factors such as *ATF4*, *MYC*, *HIF1A*, and *SREBF1* (14–17).

The transcriptional response to EAA deficiency has been studied little in the mammary glands. It has been demonstrated that the AARP is stimulated in mammary epithelial cells subjected to EAA deficiency through the eIF2α kinase GCN2 (18). Supplementing all EAA to lactating cows (19) or depleting arginine (20) produced gene expression patterns in the mammary glands consistent with the operation of the AARP. Addition of valine or methionine to cultures of mammary epithelial cells enhanced expression of the *SREBF1* product, SREBP-1c, and fat accumulation, likely through mTORC1 activation (21,22). Of the suite of potentially EAA-responsive transcription factors described above, ATF4, ATF6 and JUN regulate ER biogenesis (23–26), FOS, JUN, EGR1, MYC and HIF1A regulate cell proliferation (27–32), and SREBP-1c regulates lipogenesis (33,34). All four cellular processes are important determinants of milk yield and protein and fat content.

For this study, we hypothesized that the deprivation of Lys or Met can affect protein synthesis, ER size, cell proliferation and lipogenesis through a chronic, long-term (i.e. after 24 h) transcriptional response in bovine mammary epithelial cells (BMEC). The transcription factor genes of interest were *ATF4*, *ATF6*, *JUN*, *FOS*, *EGR1*, *MYC*, *HIF1A*, and *SREBF1*. Objectives were to evaluate if and when their expression in BMEC is affected after imposing an EAA deficiency in vitro, and to evaluate effects of EAA deficiency on protein synthesis, ER size, cell proliferation and lipogenesis.

## Materials and Methods

### Materials

All reagents and chemicals for cell culture were procured from Sigma-Aldrich (Oakville, ON, Canada) or Thermo Fisher Scientific (Waltham, MA, USA), except where noted. Kits for sample preparation and analysis were purchased from Thermo Fisher Scientific. Collagenase Type III was obtained from Worthington Biochemical Corporation (Lakewood, NJ, USA). Medium 170 was acquired from U.S. Biological (Salem, MA, USA). Radioisotopes were sourced from PerkinElmer (Waltham, MA, USA).

### Isolation and culture of bovine mammary epithelial cells

Mammary tissue was harvested from a rear quarter of lactating Holstein cows at slaughter and promptly placed in 1:1 (vol/vol) Dulbecco’s Modified Eagle Medium/Nutrient Mixture F-12 (DMEM/F12) supplemented with 1× antibiotics/antimycotics (penicillin at 100 units/ml, streptomycin at 100 µg/ml, and amphotericin B at 0.25 µg/ml). The tissue was transported on ice to the laboratory and transferred into 100-mm cell plates. Visible fat, blood, milk, and extraneous connective tissue were removed with scalpel blades. The remaining mammary tissue was minced with scalpels into approximately 1 mm^3^ fragments in a sterile environment in 5 ml of Hanks’ balanced salt solution containing 1× antibiotics/antimycotics (HBSS+). Minced tissue was transferred into a 50-ml centrifuge tube and resuspended in ice-cold HBSS+. After hand shaking the contents, the tube was placed on ice for 5 min. This procedure was repeated 4 to 5 times until the medium was clear. The washed tissue fragments were then transferred for enzymatic dissociation into sterile DMEM/F12 supplemented with 300 U/ml collagenase Type III, 1 mg/ml DNase I and 1× antibiotics/antimycotics. Digestion was carried out at 37°C with continuous shaking at 120 rpm for 4 h. Post-digestion, the preparation was filtered through a sterile sieve with 200-μm mesh and transferred into a 50-ml centrifuge tube for centrifugation at 80 × g for 30 s at 4°C. The supernatant was discarded, and the pellet of highly enriched epithelial organoids was either cryopreserved in a −80 °C freezer or seeded onto 100 mm collagen-coated dishes to allow BMEC to proliferate. If previously frozen, organoids were thawed at room temperature and then seeded onto collagen. The growth medium was composed of 1:1 (vol/vol) DMEM/F12:MCDB170, 10% fetal bovine serum, 0.1% (wt/vol) albumax II, 7.5 μg/mL bovine insulin, 0.3 μg/mL hydrocortisone, 5 ng/mL recombinant human epidermal growth factor, 2.5 μg/mL bovine apo-transferrin, 5 μM isoproterenol, 5 pM 3,3′,5-triiodo-l-thyronine, 0.5 pM β-estradiol, 0.1 nM oxytocin, and 1× antibiotics/antimycotics. 5 ml media were added every 3 d in the first week without discarding the medium in the plates and when most of the organoids were attached to the dish, then medium was replaced every 2 d. Dishes were incubated at 37°C in a humidified atmosphere of 5% CO_2_. When cells reached 50-60% confluency, they were passaged onto 100-mm cell culture plates with fresh growth medium containing 0.25% fetal bovine serum. All experiments were performed in third passage BMEC which were seeded onto 6-well plates and grown to confluency. When all wells reached confluency, cells were differentiated for 5 d in DMEM containing 3.5 mM D-glucose, 2 mM sodium acetate, 200 μM L-glutamine, and supplemented with lactogenic hormones (5 μg/mL each of bovine insulin, prolactin, and hydrocortisone), 5 μg/mL apo-transferrin, 0.5 mg/mL BSA, and 1× antibiotics/antimycotics. Media were replaced every 2 d.

### Gene expression

Differentiated BMEC were cultured in triplicate in 1 of 3 treatment media representing normal physiological concentrations of all amino acids (CTL), low Lys (LK), or low Met (LM) for 24, 40, 48, or 60 h. The base medium was DMEM containing 3.5 mM D-glucose, 2 mM sodium acetate, and 200 μM L-glutamine supplemented with lactogenic hormones (5 μg/mL each of bovine insulin, prolactin, and hydrocortisone), 5 μg/mL apo-transferrin, 0.5 mg/mL BSA, and 1× antibiotics/antimycotics. All amino acids except Met and Lys were dissolved in the base medium at their normal physiological concentrations (Mackle et al., 2000; Martineau et al., 2019) of (in µM) 180 L-alanine, 80 L-arginine, 50 L-Asparagine, 10 L-Aspartic acid, 12 L-Cysteine; 60 L-Glutamic acid, 170 L-Glutamine, 200 L-Glycine, 40 L-Histidine, 120 L-Isoleucine, 180 L-Leucine, 50 L-Phenylalanine, 70 L-Proline, 70 L-Serine, 110 L-Threonine, 30 L-Tryptophan, 50 L-Tyrosine, and 240 L-Valine. Treatments were base medium with 1) 80 μM L-Lys and 20 μM L-Met (CTL), 2) 20 μM L-Lys and 20 μM L-Met (LK), and 3) 80 μM L-Lys and 5 μM L-Met (LM). At each termination time, wells were rinsed twice and then cells were harvested into either lysis buffer from the RNA isolation kit for gene expression analysis, or RIPA buffer with protease inhibitor cocktail for protein analysis. All samples were stored at −80°C until RNA isolation and protein analysis were performed. Experiments were performed on 3 different occasions and the cells were from the same cow for each time.

Total RNA was extracted from cells with the Invitrogen PureLink RNA mini-Kit according to the manufacturer’s protocol. The RNA concentration and purity were quantified by measuring absorbance at 260 and 260/280 nm, respectively, on a NanoDrop One^C^ microvolume spectrophotometer (Thermo Fisher Scientific). All RNA integrity numbers were above 8. Isolated RNA was treated with DNase to avoid DNA contamination following the Invitrogen DNase I kit protocol. cDNA was synthesized from 1,000 ng DNase-treated RNA using the Applied Biosystems high-capacity RNA-to-cDNA kit. Primers of candidate genes were designed with the NCBI Primer-BLAST tool to yield qPCR amplification products of 80 to 150 bp (Table 1). Target genes were *ATF4*, *ATF6*, *FOS*, *JUN*, *EGR1*, *MYC*, *HIF1A*, and *SREBF1*, and reference genes were *PPIA, RPS6KB1*, and *UXT*. Primer oligos were purchased from Integrated DNA Technologies (Iowa, USA). Real-Time PCR was performed using PerfeCta SYBR Green FastMix with a StepOnePlus Real-Time PCR system (Applied Biosystems, Waltham, MA, USA). Relative gene expression was obtained by normalizing cycle threshold values for target genes to the geometric means of those for the three reference genes.

**Table 1.**
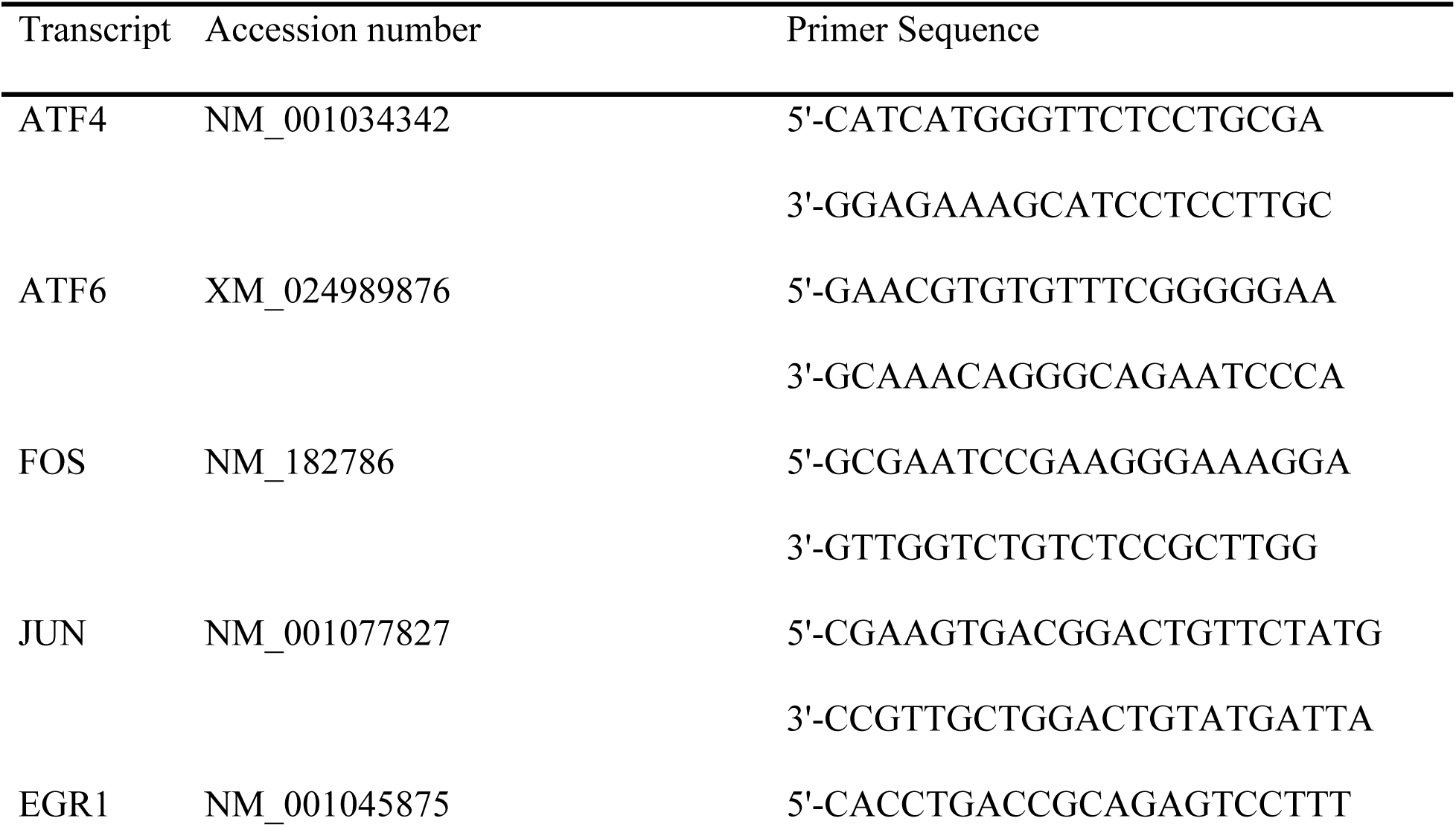
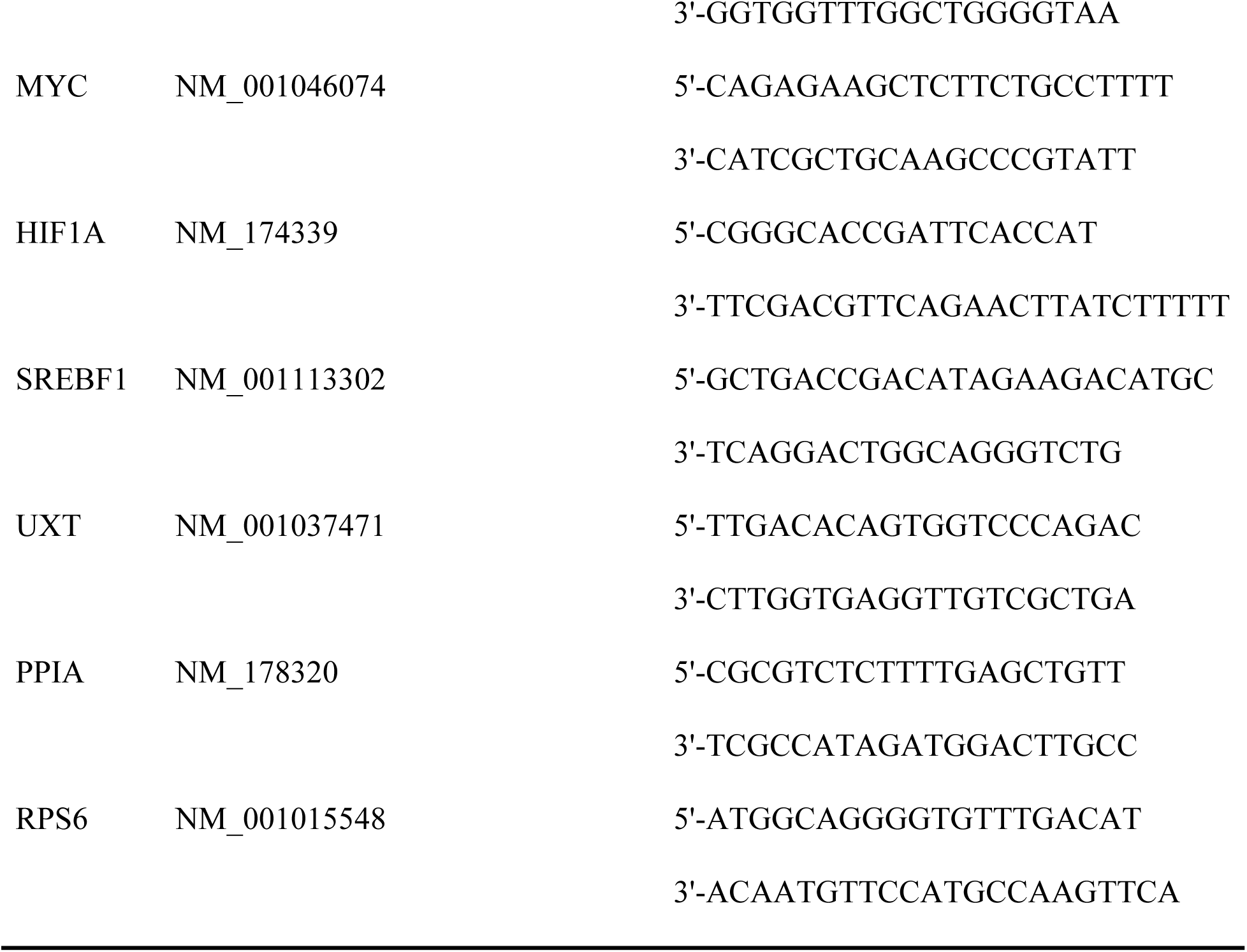
Primer sequences used in qPCR.

### Cell activity

18 wells of confluent differentiated BMEC were cultured with each of CTL, LK and LM treatment media for 48 and 60 h. Rates of protein synthesis, cell proliferation and lipogenesis were determined by quantifying, respectively, the incorporation of label from L-[2,3,4,5,6- ^3^H]phenylalanine and [methyl-^3^H]thymidine into an acid precipitate, and [1-^14^C]acetate into a hexane:isopropanol extract, over the last 2 h of the treatment period as follows. After 46 and 58 h of treatment, [^3^H]phenylalanine, [^3^H]thymidine and Na-[1-^14^C]acetate were each added to a final concentration of 1 μCi/ml into 3 different wells, for a total of 9 wells per treatment and timepoint. 2 h after isotope addition, wells were washed 3 times with PBS to remove extracellular tracer and cells were harvested into 0.5 ml RIPA lysis buffer and homogenized. Cells for protein and DNA analysis were harvested at 48 and 60 h from 9 additional wells per treatment into 0.2 ml RIPA buffer containing protease inhibitor cocktail and stored immediately at −80 °C.

DNA concentrations in 10 μl cell lysate were measured with the DNA Qubit Assay. Protein and DNA in homogenates from [^3^H]phenylalanine and [^3^H]thymidine incubations were precipitated with 0.5 ml ice-cold 20% TCA on ice for 30 min and then centrifuged at 15,000 × g for 5 min. Pellets were washed 3 times by resuspension and centrifugation with 5% TCA. After washing, pellets were dissolved in 0.5 M NaOH and 0.5 ml were transferred to 5 ml scintillation cocktail for counting in a ^3^H window. Rates of protein synthesis and cell proliferation were expressed as disintegrations per minute (DPM) per nanogram of DNA.

Lipid in lysates from [^14^C]acetate incubations was extracted into 200 ml hexane: isopropanol (3:2 vol/vol). 1 ml of the solvent layer was transferred into 5 ml scintillation cocktail and counted on a Beckman liquid scintillation counter for 10 min per sample. Lipid synthesis rate was expressed as DPM per nanogram of DNA.

### Immunoblotting

Cell proteins were extracted in RIPA buffer containing protease inhibitor cocktail and thawed on ice for 15 min. A 200-μl aliquot of the cell lysate was taken to measure protein concentration with a BCA protein assay kit using BSA as standard. Another aliquot of cell lysate was mixed with 2X Laemmli sample buffer and heated at 70°C for 10 min. 30 μg protein in cell lysates were resolved by SDS-PAGE and then transferred onto polyvinylidene difluoride membranes. Membranes were blocked in 5% (wt/vol) non-fat milk in Tris-buffered saline containing 0.1% (vol/vol) Tween 20 (TBST) at room temperature for 1 h and then incubated for 16 h with primary antibodies against the ER tracker protein disulfide isomerase (PDI; rabbit mAb #3501, Cell Signaling Technologies, Danvers, MA, USA) and a loading control GAPDH (rabbit mAb #2118) diluted in 5% non-fat milk in TBS-T at 4°C. After washing 3 times in TBS- T for 5 min, membranes were incubated with secondary antibodies diluted 1:10,000 in 5% non-fat milk in TBS-T at room temperature for 1 h with constant shaking. After washing 6 times in TBS-T, bound horseradish peroxidase-linked secondary antibodies were visualized by chemiluminescence (Bio-Rad Laboratories, Mississauga, ON, Canada). Signal intensities were quantified using Image Lab Software (Bio-Rad Laboratories) and PDI abundance was expressed relative to GAPDH.

### Statistical Analysis

Observations were subjected to analysis of variance by the GLIMMIX procedure of SAS Studio (SAS Institute Inc., Cary, NC, USA) according to the following model:

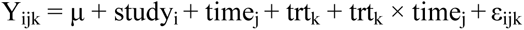

where μ is the overall mean, study_i_ is the random effect of study (i = 1 to 3), time_j_ is the fixed effect of time (j = 24, 40, 48, 60, or j = 48, 60), trt_k_ is the fixed effect of treatment (k = 1 to 4), and ε_ijk_ is random variation. Effects of treatments within and across timepoints were estimated as linear contrasts between CTL, LK, and LM. Contrasts were considered statistically significant when P ≤ 0.05 and trends when 0.05 < P ≤ 0.15.

## Results

### Gene expression

To evaluate if and when the selected transcription factors *ATF4*, *ATF6*, *JUN*, *FOS*, *EGR1*, *MYC*, *HIF1A* and *SREBF1* were affected by Lys or Met deprivation, their mRNA expression was measured at 24, 40, 48 and 60 h after Lys or Met subtraction to ¼ the normal plasma concentration.

Compared to CTL, LK and LM both increased *ATF4* expression (*P* < 0.05; Table 2), particularly at 40 h (*P* < 0.01; Figure 1). LM increased *ATF6* expression across all timepoints (*P* = 0.012; Table 2) while LK did not (*P* = 0.77). LM tended to increase *ATF6* expression at 40 (*P* = 0.08) and 60 h (*P* = 0.11) compared to CTL (Figure 1). The expression of *JUN* tended to increase with LM (*P* = 0.08) but was not affected by LK (*P* = 0.17). At 60 h, both LK and LM exhibited lower expression of *JUN* (*P* < 0.01) compared to CTL (Figure 1) which appears to have been due to an increase in *JUN* expression in CTL incubations that did not occur on LK or LM. There was no significant effect or trend for the expression of *FOS* or *MYC* on either LK or LM (Table 2; Figure 1). The expression of *EGR1* was decreased by LM (*P* = 0.04; Table 2) but there was no significant effect at any single timepoint for either of the treatments (Figure 1). LM increased the expression of *HIF1A* across all timepoints (*P* < 0.01; Table 2), and at 24 h (*P* < 0.01; Figure 1), and tended to increase *HIF1A* expression at 48 h (*P* = 0.12). *SREBF1* expression was decreased by LK (*P* = 0.05) but not by LM. LK decreased *SREBF1* expression at 24 h (*P* = 0.01; Figure 1) while LM increased *SREBF1* expression at 40 h (*P* = 0.02) and decreased its expression at 48 and 60 h (*P* = 0.01).

**Figure 1.**
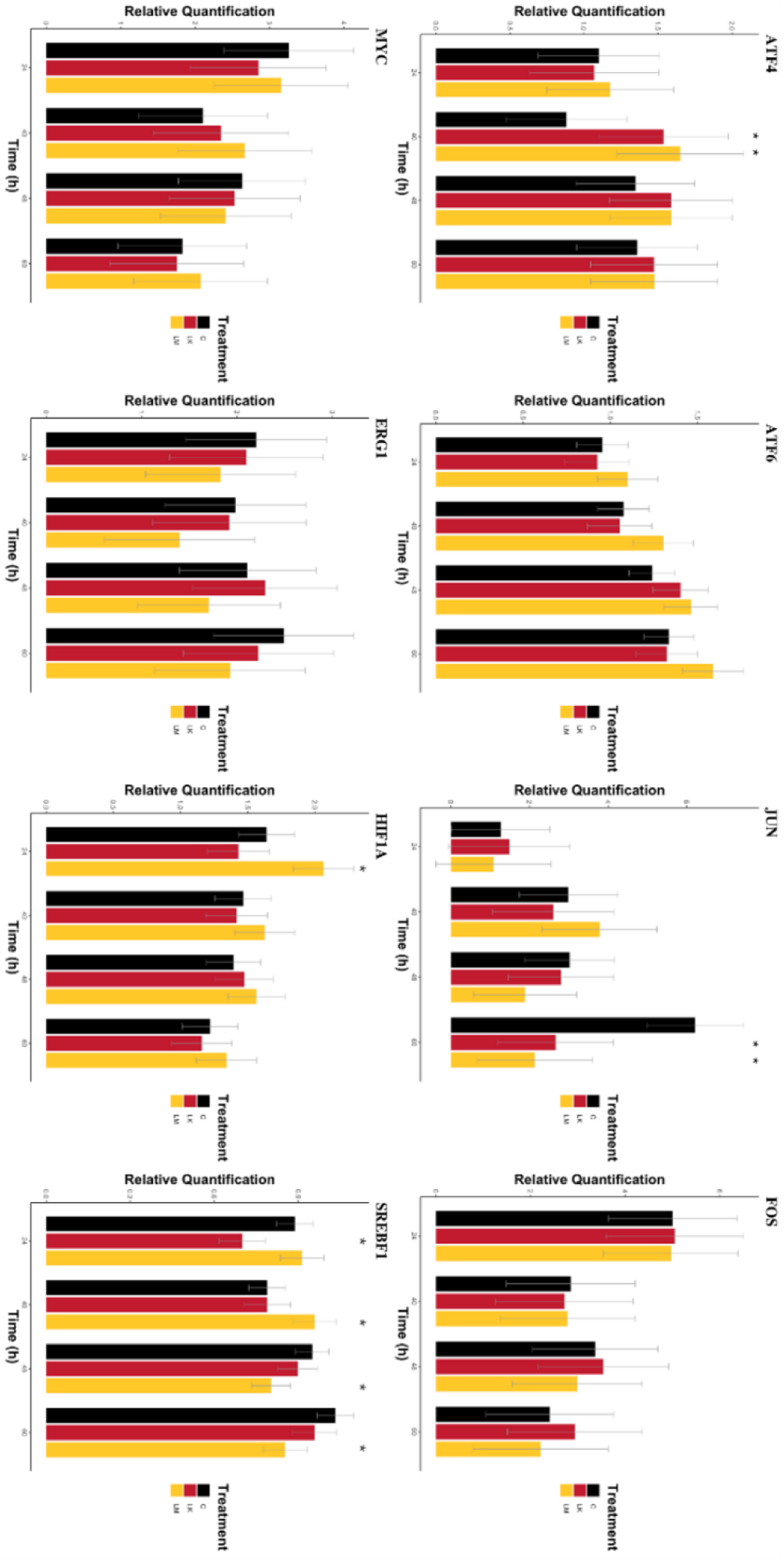
Gene expression of candidate transcription factors in bovine mammary epithelial cells at 24, 40, 48, 60 h after lysine or methionine subtraction to ¼ the normal concentration. Values are least square means ± SE (n = 3) in arbitrary units. CTL = control; LK = low lysine; LM = low methionine. \**P* ≤ 0.05, †0.05 < *P* ≤ 0.15 compared to CTL.

**Table 2.** Gene expression of candidate transcription factors in bovine mammary epithelial cells across all timepoints of 24, 40, 48 and 60 h after lysine or methionine limitation to ¼ the normal concentration^1^.

| Gene | Treatment<sup>2</sup> |  |  | SEM | P-value |  |
| --- | --- | --- | --- | --- | --- | --- |
|  | CTL | LK | LM |  | LK | LM |
| ATF4 | 1.17 | 1.42 | 1.47 | 0.40 | 0.04 | 0.01 |
| ATF6 | 1.15 | 1.18 | 1.36 | 0.13 | 0.77 | 0.01 |
| FOS | 3.41 | 3.56 | 3.24 | 1.30 | 0.71 | 0.65 |
| JUN | 3.37 | 2.39 | 2.22 | 1.10 | 0.17 | 0.09 |
| EGR1 | 2.20 | 2.13 | 1.72 | 0.71 | 0.81 | 0.05 |
| MYC | 2.46 | 2.37 | 2.58 | 0.85 | 0.69 | 0.54 |
| HIF1A | 1.43 | 1.37 | 1.65 | 0.20 | 0.43 | <0.01 |
| SREBF1 | 0.91 | 0.84 | 0.88 | 0.06 | 0.05 | 0.36 |

<sup>1</sup>Values are least squares means ± SE (n = 12) in arbitrary units.
<sup>2</sup>CTL = control; LK = low lysine; LM = low methionine.

### Cell activity

As presented in Figure 2, LK decreased protein synthesis 21% at 48 h (*P* = 0.04) and 22% at 60 h (*P* = 0.02), while LM decreased protein synthesis 34% at 48 h (*P* < 0.01) and 29% at 60 h (*P* < 0.01). Neither LK nor LM had any effects on fat synthesis rates. LK decreased cell proliferation 26% at 48 h (*P* = 0.03) but showed no effect at 60 h. LM dropped cell proliferation 48% at 48 h (*P* < 0.01) and 22.2% at 60 h (*P* = 0.01). PDI expression was measured as an ER abundance marker. There were no effects of LK or LM on PDI expression at either 48 h or 60 h.

**Figure 2.**
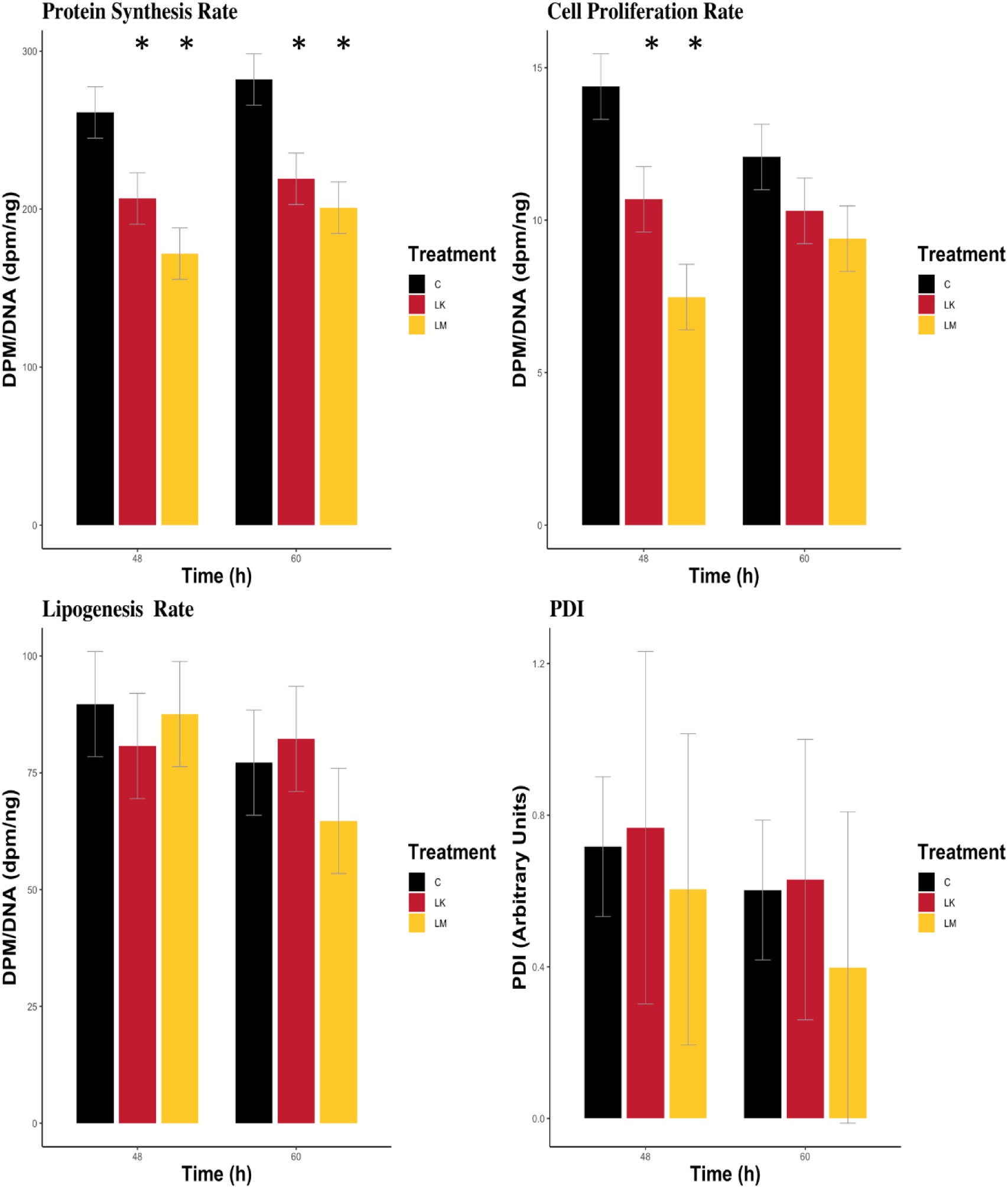
Cell activities in bovine mammary epithelial cells across all timepoints of 48 and 60 h after lysine or methionine limitation to ¼ the normal concentration. Values are least square means ± SE (n = 3). CTL = control; LK = low lysine; LM = low methionine. \**P* ≤ 0.05 compared to CTL.

**Table 3.** Cell activities in bovine mammary epithelial cells across all timepoints of 48 and 60 h after lysine or methionine limitation to ¼ the normal concentration^1^.

| Item | Treatment<sup>2</sup> |  |  | SEM | P-value |  |
| --- | --- | --- | --- | --- | --- | --- |
|  | CTL | LK | LM |  | LK | LM |
| Protein synthesis rate | 272 | 213 | 186 | 11.5 | <0.01 | <0.01 |
| Fat synthesis rate | 83.4 | 81.5 | 76.1 | 8.0 | 0.87 | 0.53 |
| Cell proliferation rate | 13.2 | 10.5 | 8.4 | 0.8 | 0.03 | <0.01 |
| PDI (arbitrary units) | 0.66 | 0.70 | 0.50 | 0.03 | 0.93 | 0.72 |

<sup>1</sup>Values are least squares means ± SE (n = 6) in dpm/ng DNA unless otherwise stated.
<sup>2</sup>CTL = control; LK = low lysine; LM = low methionine; PDI = endoplasmic reticulum tracker protein disulfide isomerase.

## Discussion

To elucidate the effects of reduced concentrations of a single EAA on BMEC activity and expression of genes for 8 transcription factors hypothesized to be regulated by EAA following different signaling mechanisms and affecting different downstream activities related to milk synthesis, experiments were performed with BMEC in culture. Treatments were designed to provide Lys and Met concentrations at ¼ of normal physiological concentrations in plasma of lactating dairy cows while maintaining all other amino acids at normal levels. Measurements were made after 24 h to focus on longer term responses to chronic EAA deficiency. While changes in *HIF1A* and *SREBF1* expression were detected at 24 h, changes in expression of *ATF4*, *ATF6*, *EGR1*, *HIF1A* and *SREBF1* occurred after 40 h. Similarly, others have reported that expression of *ATF4* and its downstream targets is not upregulated in response to deprivation of a single amino acid until more than 12 h of incubation (18,35) and can remain elevated for 72 h (36,37). *FOS* and *JUN* are considered immediate-early genes and have been found to peak in expression around 6 h into an EAA deficiency and return to normal by 24 h (11), which is consistent with our finding of no effect after 24 h. *EGR1* expression can also constitute part of the early response to stress (12) but Papež et al. (37) found no effect of glutamine deprivation on *EGR1* expression in CHO cells until 48 h, when it declined. *HIF1A* expression was also downregulated in response to glutamine deprivation at 48 h of incubation (37). Because most of the transcription factor changes occurred after 40 h, we chose 48 and 60 h as timepoints for cell activity measurements.

Deficiencies of both Lys and Met suppressed protein synthesis and activated *ATF4* expression, indicating the classic AARP had been triggered, likely through free tRNA-mediated activation of GCN2 (18). ATF4 homo- and hetero-dimerizes with ATF6, FOS and JUN, among other basic leucine-zipper transcription factors, to induce expression of genes involved in resolving and adapting to the EAA deficiency, such as amino acid transporters and ER-resident proteins (8). In addition to being up-regulated when global protein synthesis is inhibited by eIF2α phosphorylation, ATF4 and its partners can also induce expression of activators and inhibitors of protein synthesis, possibly to affect how the system responds to other regulatory influences via mTORC1, such as growth factors and ATP concentration (17). Interestingly, muscle-specific knockout of *ATF4* promoted expression of anabolic genes involved in protein synthesis and prevented the age-related decline in muscle mass (38). Thus, the upregulation of *ATF4* expression we observed during EAA deficiency may have contributed to the decreased rate of protein synthesis.

Despite a decrease in global protein synthesis, there was no effect of Lys or Met deficiency on ER size indicated by PDI abundance. This finding may indicate that proteostasis was achieved by *ATF4* up-regulation. *ATF6* is itself an ATF4 target (9) whose gene product can be activated by the ISR to selectively increase the expression of genes related to ER biogenesis and mRNA translation (39). A mix of branched-chain amino acids dose-dependently increased *ATF6* expression in mouse cardiac myocytes (40), and EAA activated the ATF6 target XBP1 in the mammary glands of cows to allow for greater milk protein yields (19). The increase in *ATF6* expression we detected during Met deficiency may have contributed to maintenance of the capacity for synthesis of the secretory proteins of milk.

Expression of *JUN* and *FOS* to produce hetero-dimerization partners of ATF4 is considered part of an early stress response through RAS/RAF/MEK/ERK signaling that promotes cell survival by stimulating proliferation and suppressing apoptosis (41). Exposing cells to an absence of a single EAA, or to histidinol to block histidyl-tRNA synthesis, caused increases in *JUN* and *FOS* expression within 12 h (11,12,42,43). In our experiments, after 24 h, there were no differences in *JUN* or *FOS* expression besides a lack of increase in *JUN* at 72 h. The findings suggest *JUN* and *FOS* do not participate in the long-term signaling responses in BMEC.

As its name suggests, *EGR1* is also categorized as an early response gene and, like *FOS* and *JUN*, its expression is activated by the RAS/RAF/MEK/ERK signaling pathway (29). Supplementation with histidinol or induction of ER stress with thapsigargin increased the expression of *EGR1* in HepG2 cells from 8 to 24 h of incubation (12,13). Contrary to these findings, our data demonstrate that Met deficiency led to a long-term suppression of *EGR1* expression. Glutamine deficiency in CHO cells also led to a long-term decrease in *EGR1* expression at 48 h without a prior early response (37). *EGR1* is a noted driver of cell proliferation (44) and its downregulation may have contributed to the decrease in cell proliferation we observed in BMEC exposed to LM.

Another stress-induced regulator of cell proliferation is HIF1A, which is responsible for cell cycle arrest during hypoxia (27,45). The arrest mechanism involves antagonism by HIF1A of the proliferative effects of the transcription factor MYC (45,46), and it has been proposed that the balance between HIF1A and MYC influences the decision to progress through the cell cycle. The increased expression of *HIF1A* during Met deficiency in our experiment, accompanied by no change in *MYC* expression, may have contributed to the observed decrease in cell proliferation rate. We can only speculate as to how EAA deficiency may provoke *HIF1A* and *MYC* expression. *MYC* expression can be upregulated by the same RAS/RAF/MEK/ERK signaling pathway responsible for *JUN*, *FOS* and *EGR1* up-regulation (47), which we did not observe. Alternatively, expression of both transcription factors can be stimulated by mTORC1 signalling (14,15) which is depressed by EAA deficiency (2,48). However, removal of any EAA from CHO cell cultures increased the abundance of *MYC* mRNA within 1 to 8 h (43), while removal of glutamine increased *MYC* expression and decreased *HIF1A* expression after 48 h of incubation (37). Further research may help clarify the relationship between EAA deficiency and the *HIF1A*/*MYC* balance.

SREBP-1c is the key lipid biosynthesis transcription factor in various tissues (33,34), and is a major regulator of milk fat synthesis in BMEC (49,50). The expression of *SREBF1* and its lipogenic targets is diminished by inhibiting mTORC1 (14,50,51), which EAA can regulate directly in mammary cells (2). Indeed, valine addition to porcine mammary epithelial cell cultures increased nuclear SREBP-1c and intracellular triacylglycerol contents in an mTORC1- dependent manner (22), and methionine addition to supraphysiological concentrations for BMEC enhanced SREBP-1c abundance and fat accumulation (21). In our experiments, Met and Lys deficiency failed to affect lipogenesis rates at 48 and 60 h even though LM decreased the expression of *SREBF1* at those timepoints. However, LM also increased *SREBF1* expression at 40 h and this inconsistent effect may have clouded the lipogenic response.

## Conclusions

In this study, EAA deficiency decreased the rate of protein synthesis and increased the expression of *ATF4* which indicates the classic AARP was triggered in BMEC. The alternative RAS/RAF/MEK/ERK stress signaling pathway did not appear to be an important player in the long-term response to EAA deficiency as evidenced by a lack of increased expression of *JUN*, *FOS* or *EGR1* between 24 and 60 h. The lower protein synthesis rate can affect cell proliferation rates but the decreased expression of *EGR1* and increased expression of *HIF1A* during Met deficiency may have also contributed to the reduced cell proliferation rates. These results suggest that a transcriptional response to EAA deficiency contributes to effects on protein synthesis and cell proliferation in BMEC.

## Acknowledgements

The authors wish to thank Julie Kim, Sara Ahmady, Jillian Wang, and Brian MacDougall at the University of Guelph for technical assistance.

